# Targeted and passive environmental DNA approaches outperform established methods for detection of quagga mussels, *Dreissena rostriformis bugensis* in flowing water

**DOI:** 10.1101/2020.01.07.890731

**Authors:** Rosetta C Blackman, Kar Keun Sean Ling, Lynsey R Harper, Peter Shum, Bernd Hänfling, Lori Lawson Handley

## Abstract

1. The early detection of invasive non-native species (INNS) is important for informing management actions. Established monitoring methods require the collection or observation of specimens, which is unlikely at the beginning of an invasion when densities are likely to be low. Environmental DNA (eDNA) analysis is a highly promising technique for the detection of INNS – particularly during the early stages of an invasion.
2. Here, we compared the use of traditional kick-net sampling with two eDNA approaches (targeted detection using both conventional and quantitative PCR, and passive detection via metabarcoding with conserved primers) for detection of quagga mussel, *Dreissena rostriformis bugensis;* a high priority INNS, along a density gradient on the River Wraysbury, UK.
3. All three molecular tools outperformed traditional sampling in terms of detection. Conventional PCR and qPCR both had 100% detection rate in all samples, and outperformed metabarcoding when the target species was at low densities. Additionally, quagga mussel DNA copy number (qPCR) and relative read count (metabarcoding) were significantly influenced by both mussel density and distance from source population, with distance being the most significant predictor.
4. *Synthesis and application*. All three molecular approaches were more sensitive than traditional kick-net sampling for the detection of the quagga mussel in flowing water, and both qPCR and metabarcoding enabled estimates of relative abundance. Targeted approaches were more sensitive than metabarcoding, but metabarcoding has the advantage of providing information on the wider community, and consequently impacts of INNS.

## Introduction

Invasive non-native species (INNS) have wide ranging effects on ecosystems, including biodiversity loss (Doherty et al., 2016; Gallardo et al., 2016) and economic impacts including damage to infrastructure (Pimentel et al., 2005; Connelly et al., 2007). Eradication or containment methods are most effective during the early stages of an invasion, both in terms of success and financial cost (Hulme, 2006; Mehta et al., 2007). Early detection is particularly challenging for species that are small, elusive or cryptic and these species often go unnoticed until they become established (Simmons et al., 2015; Blackman et al., 2017). Monitoring organisms such as freshwater invertebrates generally relies on specimen collection and morphological identification, and newly invading species can be easily overlooked. Alternative methods that offer rapid and cost-effective detection of new INNS are crucial at a time when rapid increase in trade and transport has caused a surge in the number of successful invasions to new environments (Hulme, 2006).

The use of molecular methods to identify DNA taken from environmental samples (i.e. environmental DNA or “eDNA”) has been a major research focus in biodiversity monitoring over the last 10 years, including the detection of INNS (Ficetola et al., 2008, Thomsen et al., 2012, Blackman et al., 2018). Two broad approaches can be applied to detect a species of interest. The “targeted” approach utilises species-specific primers with conventional PCR (cPCR), quantitative PCR (qPCR) or droplet digital PCR (ddPCR) to detect a single species. By contrast, metabarcoding can be considered a “passive approach” as it does not focus on single species (Lawson Handley, 2015; Creer et al., 2016; Deiner et al., 2017). Metabarcoding utilises conserved primers to amplify DNA from a group of taxa e.g. fish and other vertebrates (e.g. Kelly et al., 2014; Miya et al., 2015; Simmons et al., 2015), zooplankton (e.g. Brown et al., 2016), or insects (e.g. Yu et al., 2012., Elbrecht et al., 2016). Amplicons are then sequenced using High Throughput Sequencing (HTS) platforms and bioinformatically assigned to taxa. Metabarcoding provides an opportunity to detect multiple species simultaneously, including those, which have not been listed as priority INNS (Simmons et al., 2015; Blackman et al., 2017). Comparisons between traditional methods for biodiversity monitoring and targeted (Dejean et al., 2012; Tréguier et al., 2014) or passive (Smart et al., 2015; Hänfling et al., 2016; Wilcox et al., 2016) molecular approaches have demonstrated an increased probability of detection using eDNA methods.

The targeted approach was first used by Ficetola et al. (2008) to detect invasive American bullfrog *Lithobates catesbeianus*, in lentic waterbodies and has since been applied to over 100 target INNS (Blackman et al., 2018). Conventional PCR is a well-established and widely practiced method for detecting presence/absence of target DNA by visualisation of PCR products on agarose gels. However, quantitative PCR (qPCR) has on many occasions been shown to be more sensitive than PCR (e.g. Nathan et al., 2014; Simmons et al., 2015; De Ventura et al., 2016) and therefore more suitable for detecting INNS at low density, e.g. the early stages of invasion. Quantitative PCR has the added advantage of providing the number of DNA copies in a sample, which has been successfully linked to biomass or abundance of some taxa (e.g. Amphibians, Thomsen et al., 2012; Fish, Takahara et al., 2012; Gastropods, Goldberg et al., 2013). However, qPCR requires the construction of calibration curves from known standard concentrations and is substantially more expensive and time-consuming than cPCR (Nathan et al., 2014) and maybe less accessible to end-users than cPCR (Blackman et al., 2020).

The potential of eDNA metabarcoding for INNS monitoring has already been shown as an effective tool (Simmons et al., 2015; Brown et al., 2016; Blackman et al., 2017; Borrell et al., 2017; Holman et al., 2019). These studies include the detection of new INNS, which had been previously overlooked by traditional methods. For example, Blackman et al. (2017) used a metabarcoding approach to investigate macroinvertebrate diversity and discovered a non-native Gammaridae species, *Gammarus fossarum*, which traditional methods had failed to identify for over 50 years in the UK. Similarly, Simmons et al. (2015) detected the Northern Snakeshead, *Channa argus*, in the Muskingum River Watershed, Ohio, which had not been included on INNS priority lists. Metabarcoding also has the potential to act as a surveillance tool and has been applied to monitor high-risk pathways, such as ballast water (Ardura et al., 2015; Zaiko et al., 2015), the live bait trade (Mahon et al., 2014) and ports (Brown et al., 2016; Borrell et al., 2017). However, comparisons of targeted and passive approaches for INNS or other focal species have been exemplified in the bighead carp, *Hypophthalmichthys nobilis* (Simmons et al., 2015) and great crested newts, *Triturus cristatus* (Harper et al., 2018). These studies both showed that a targeted approach had a higher sensitivity than the passive approach. Possible causes of this include primer choice and PCR bias, i.e. species successfully amplifying and masking others in the PCR reaction during metabarcoding (Harper et al., 2018). Further comparisons of these methods covering different target taxa are required to better inform management strategies.

In this study we focus on the quagga mussel, *Dreissena rostriformis bugensis*, a highly invasive bivalve from the Ponto-Caspian region, which has a significant impact on all trophic levels within the environments it invades (Karatayev et al., 2007; Roy et al., 2014). In a horizon scanning exercise from 2014, the quagga mussel scored the highest out of 591 species evaluated for its potential to invade, establish and have negative impact in the UK (Roy et al., 2014). Consistent with predictions, quagga mussels were subsequently detected later the same year (Aldridge et al., 2014). Due to its wider tolerance to environmental conditions, it has the potential to spread rapidly, and occupy a greater range of habitat than the closely related zebra mussel, *Dreissena polymorpha*, which has already spread throughout much of Europe and North America (Nalepa et al., 2010; Gallardo and Aldridge 2013; Quinn et al., 2014). Motivated by a need for cost effective tools for quagga mussel detection, we previously designed a cPCR assay, which provided 100% detection rate in field trials with quagga mussel populations (Blackman et al., 2020). Here we further developed and validated this assay for abundance estimation using a dye-based qPCR approach in mesocosm experiments and field trials. Secondly, we directly compared the three molecular methods (cPCR, qPCR and metabarcoding) to traditional kick-net sampling for detection of quagga mussels in the field. Finally, we investigated whether qPCR and metabarcoding are suitable for estimating relative abundance of quagga mussels by sampling a density gradient away from the main population. We hypothesized that: (1) the probability of detecting quagga mussels will be higher for eDNA methods than traditional methods, (2) targeted methods will have a higher probability of detecting quagga mussel than the passive method, (3) qPCR will be more sensitive than cPCR for quagga mussel detection and (4) both read count and DNA copy number will decline with distance downstream of the main population.

## Methods

### Field sample collection, DNA capture and extraction

Quagga mussels were first detected in the UK in Wraysbury Reservoir, Berkshire, in October 2014. This site is considered the founding population of quagga mussels in the UK. The mussels spread through the outfall and along the adjacent River Wraysbury, which is a tributary of the River Colne. The species currently has a restricted distribution in the South East of England but has also been recorded in reservoirs adjacent to the River Lee, north of London, and in the River Thames. Samples were collected from Wraysbury Reservoir and River in May 2016, working in an upstream direction to ensure DNA availability and inhibitor levels were not altered by sampling activity. In order to compare established and molecular methods directly, we collected eDNA and kick-net samples at six sites across the river at each of three main sampling locations downstream of Wraysbury Reservoir: Wraysbury Weir (WW, 0.61 km downstream of Wraysbury Reservoir), Wraysbury Bridge (WB, 1.10 km), and Wraysbury Gardens (WG, 2.7 km) (orange diamonds, Fig. 1 and Supplementary Information I Table S1). Water samples from these three locations consisted of three 500 ml replicates, collected at each of six sites across the width of the river (n = 54). eDNA samples were collected from the river surface, using sterile gloves and Gosselin™ HDPE plastic bottles (Fisher Scientific UK Ltd, UK) prior to kick-net sampling. Three-minute kick-net samples were collected from the same six sites at each of these three locations, in accordance with established methods (Murray-Bligh, 1999). eDNA sampling of three 500 ml replicates was also carried out at four additional locations along Wraysbury River (Reservoir Outfall, RO, 0.2 km; Moor Lane, ML, 1.7 km; Hale Street, HS, 2.72 km; and Upper Thames Confluence, UT, 2.75 km, blue diamonds, Fig. 1, Supplementary Information I Table S1). These additional eDNA samples were collected to provide additional fine-scale resolution in the eDNA analysis, and were analysed with PCR and qPCR only, whereas samples from the three main locations were analysed with PCR, qPCR and metabarcoding. Sampling locations within the river were chosen based primarily on accessibility, in accordance with standard macroinvertebrate monitoring procedures carried out by UK agencies. We did not target specific locations with previous records of quagga mussels, or their preferred habitat. At the time of sampling, quagga mussels had not been reported upstream of the reservoir. We therefore collected an eDNA sample upstream of the reservoir with the expectation that this would be negative for quagga mussel (Waterside Car Park, n = 3). Sample bottles filled with ddH2O were also taken into the field as filtering blanks (n = 3). Hence the total number of eDNA samples collected in the field was n = 72. All eDNA samples and blanks were stored in a cool box with ice packs and transported to the laboratory for filtration within 24 hours. Water samples were filtered through 0.45 μm cellulose nitrate membrane filters attached to a vacuum pump and DNA extractions performed following Brolaski et al. (2008) with minor modifications (see Supplementary Information I and II for full details). Kick-net samples were placed in individual 42 fluid oz. sterile Whirl-pak^®^ bags (Cole-Palmer, Hanwell, London) and stored in a cool box before being frozen at −20°C in the laboratory. Each sample was thawed at room temperature and rinsed with ddH2O separately prior to analysis. Samples were then sorted for the presence of *D. r. bugensis* only and all specimens were counted and subsequently stored at −20°C.

**Figure 1:**
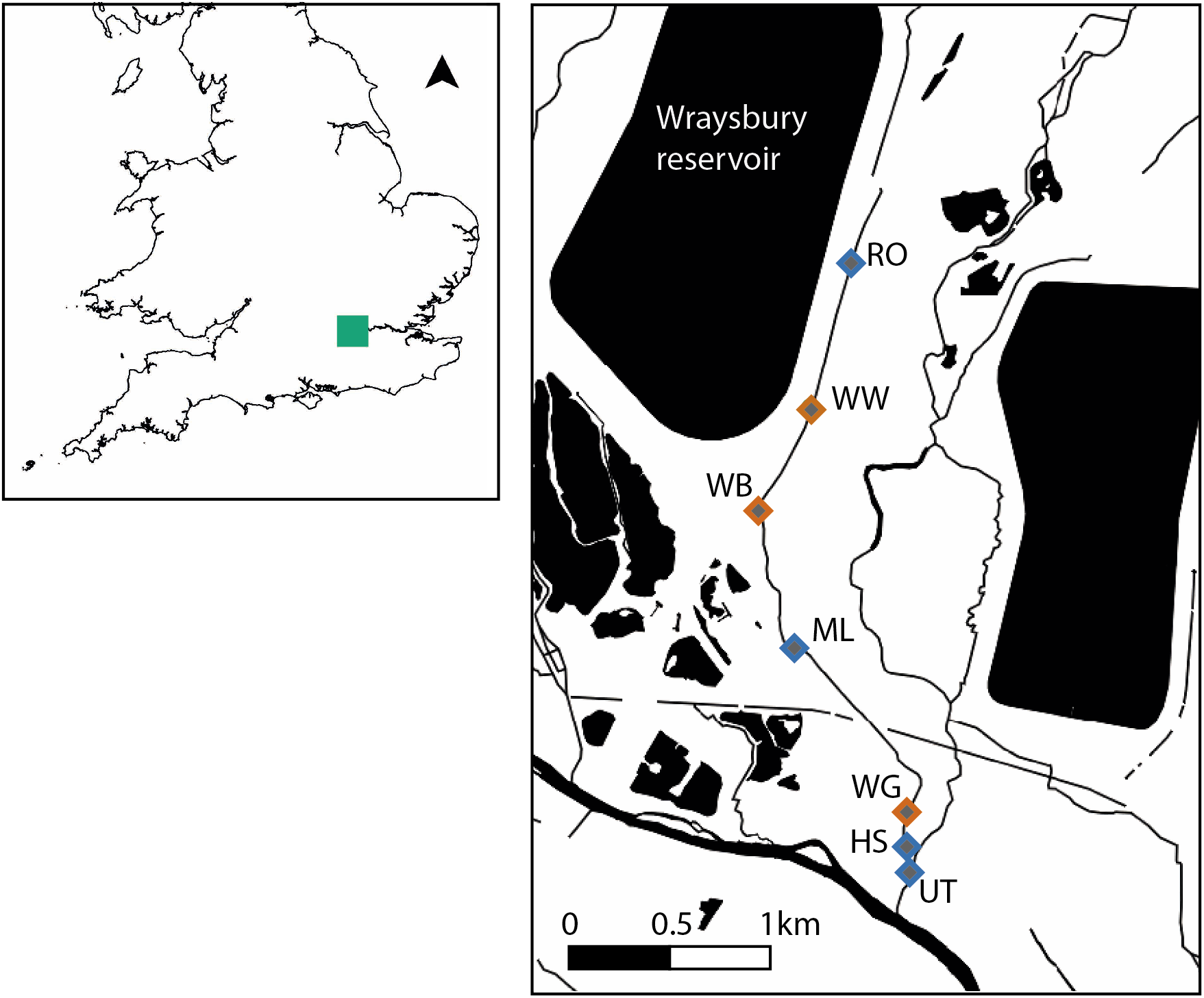
Sampling locations on the Wraysbury River. Diamond symbols represent the sampling locations: Reservoir Outfall (RO, 0.2 km downstream of the reservoir), Wraysbury Weir (WW, 0.61 km), Wraysbury Bridge (WB, 1.10 km), Moor Lane (ML, 1.7 km), Wraysbury Gardens (WG, 2.7 km), Hale Street (HS, 2.72 km), and Upstream Thames confluence (UT, 2.75 km). eDNA and kick-net samples were collected from sites with an orange outline and sites sampled for eDNA only have a blue outlined symbol (Full grid references are provided in Supplementary Information I Table S1).

### Performance of targeted molecular detection

The performance of targeted DNA assays for the detection of invasive quagga mussel DNA was first examined through laboratory-based mesocosm experiments. The objectives of these experiments were a) to validate species-specific DNA assays, using both conventional and quantitative PCR, to target eDNA detection of quagga mussels; b) to assess the duration and persistence of eDNA signal over a 42-day period; and c) to examine a correlation between eDNA concentration and known densities of quagga mussel. The mesocosm experiment was set up to monitor DNA production of mussels at three densities (1, 5 and 20 individuals, with total and individual biomass weights recorded) over 21 days, with three replicates of each density and a control tank with no mussels present. Fifteen litre plastic tanks with individual aeration stones were set up in a climate controlled facility where temperature averaged 16 °C (range 14–18 °C), with light:dark cycles of 16 h:8 h. Sampling events took place over 42 days at: 0hrs (prior to specimens being added), 4 hrs, 8 hrs, 24 hrs, 48 hrs, 72 hrs, 7 days, 15 days and 21 days. On day 21, the specimens were removed, and sampling continued at: 22 days, 28 days, 35 days, 42 days (n = 130). Two hundred millilitre water samples were collected at each time point and each sample was filtered through a single filter membrane following the same protocol as the field samples (Blackman et al., 2020).

#### Conventional PCR (cPCR) assay

We used a cPCR assay to detect the quagga mussel, *D. r. bugensis*, using a species-specific primer pair targeting a 188 bp fragment of the cytochrome oxidase I (COI) gene (DRB1_F – 5’-GGAAACTGGTTGGTCCCGAT3’- and DRB1_R 5’-GGCCCTGAATGCCCCATAAT-3’, see Supplementary Material I for full PCR protocol). The DRB1 PCR assay has shown to be highly sensitive (positive detection with 0.001 ng of target tissue DNA per reaction) and does not cross amplify closely related Dreissenid or native mussel species (Blackman et al., 2020). All mesocosm (n=130), field samples (n = 66) and control samples (field (3), filter (3) and PCR (6) blanks) were tested with the DRB1 assay. A tissue DNA sample from closely related zebra mussel was included as an additional negative control.

#### Quantitative PCR (qPCR) assay

The DRB1 assay was further developed using fluorescent dye-based qPCR (PowerUp SYBR green, Thermo Fisher, UK, see Supplementary Material I for full details). Briefly, we performed serial dilutions of DNA from tissue samples (from 1 to 1 x 10^−7^ ng/μl) to estimate the LoD, and performed the DRB1 qPCR assay on samples collected from the mesocosm experiment, field an control samples. Zebra mussel tissue DNA was used as a negative control in the qPCR assays. Amplification curves, Cq values and melt curves were analysed using StepOne-PlusTM software 2.0. Quantification calibration curves using a synthesized gBlock of the target fragment was used to estimate the DNA copy number for each sample (See supplementary Information I and IV for full details).

### Passive detection: eDNA metabarcoding

Degenerate metazoan primers that target the conserved COI region were used for metabarcoding (Geller et al., 2013 and Leray et al., 2013): jgHCO2198: TAIACYTCIGGRTGICC RAARAAYCA and mICOIintF: GGWACWGGWTGAACWGTWTAYCCYCC. These primers were chosen as they i) target the same mitochondrial gene as our targeted assays, ii) are the most commonly used COI metabarcoding primers and are good candidates for studying the entire river community, and iii) we confirmed in previous work that they amplify Dreissenid mussels. We used universal primers rather than mollusc specific metabarcoding primers (which are also available, Klymus et al., 2017), since universal primers are more likely to be useful in routine community monitoring of INNS. We confirmed amplification of the target taxa from tissue extractions prior to library preparation. Full details of the library preparation are provided in Supplementary Information I. In brief, the 54 field samples from WW, WB and WG (Fig. 1) were sequenced, with the addition of 3 field blanks, 3 filter blanks, 6 PCR blanks and 3 positive non-target tissue samples (2 each of *Triops cancriformis and Osmia bicornis total n* = 69). A two-step PCR protocol was used for library preparation: first the target region was amplified for each sample and a second amplification was performed to add the Illumina adaptor sequences onto the initial amplicons. Samples were normalized following the second PCR using SequalPrep Normalization plates (Invitrogen, UK) and samples were then pooled in equimolar concentrations. The final pooled library was concentrated, cleaned with a QIAquick Gel Extraction Kit (Qiagen, UK), quantified and sequenced at 13 pM concentration with 20% PhiX, on an Illumina MiSeq platform using 2 x 250bp v2 chemistry at the University of Hull.

### Bioinformatics and data analysis

Full details of the bioinformatics pipeline are provided in Supplementary Material I. In brief, processing of Illumina data and taxonomic assignment were performed using a custom pipeline, MetaBEAT (metaBarcoding and eDNA Analysis Tool) v 0.8 (https://github.com/HullUnibioinformatics/metaBEAT) but with minor modifications including filtering for the presence of the target taxa, *D. r. bugensis*, by comparing against curated reference sequences from GenBank, listed in Supplementary Material I.

We normalised the metabarcoding data to account for uneven sequencing depth between samples, by dividing the number of reads for each OTU within a sample by the total number of reads in that sample and multiplying by 100 (hereafter referred to as relative read count).We used Generalized Linear Models (GLMs) to investigate 1) the influence of mussel density, biomass and time on the DNA copy number (qPCR) since the start of the mesocosm experiment (before the mussels were removed from the tanks only), and 2) the influence of mussel density and distance from source population on the DNA copy number (qPCR) from the field samples and relative read count (metabarcoding). Initial model tests showed all data sets were overdispersed, therefore a negative binomial GLM with a log link function was used to resolve this issue. To check for model suitability, we checked the residual deviance of each model fitted a chi square distribution (see Supplementary Information III Tables S2, S3 and S4 for GLM results). Finally, we assessed the relationship between DNA copy number and relative read count with the distance along the river using all seven sites included in the qPCR experiment, using a Spearman’s Rank Correlation since the data were not normally distributed. All calculations and visualizations of the data were carried out in R (R-Core-Team 2017), with GLMs performed using the MASS package (Venables et al., 2002).

## Results

We successfully adapted the DRB1 primer pair for detection of quagga mussel with the addition of SYBR green. Across both traditional and eDNA field experiments we were able to detect quagga mussel with kick-net sampling, cPCR, qPCR and metabarcoding. All negative controls (field blanks, filtration blanks, PCR blanks and non-target tissue blanks were negative for cPCR and qPCR. Minor contamination was detected in 3 field blanks with metabarcoding, with low reads of *Homo sapiens (5 and 14 reads), Triops cancriformis* (the positive tissue extract) (10 reads) and unassigned reads (42-189 reads). We therefore applied a 0.2% read threshold to remove possible contamination from the metabarcoding dataset (i.e. sequences that had a frequency of 0.2% or less in each sample were removed).

### qPCR assay development and validation in mesocosms

The limit of detection (LoD) determined from the serial dilutions of tissue DNA was 1 x 10^−4^ ng/μl per reaction with both cPCR and qPCR, which equates to approximately 5 copies per reaction (See Supplementary Information III Fig. S1 and Fig. S2 for amplification plots). Rapid DNA accumulation in the first two days of the mesocosm experiments was followed by a depletion in DNA concentration until removal of mussels on day 21 of the experiment (Fig. 2). This pattern of DNA accumulation and depletion is highly consistent across all three density treatments. Time since the beginning of the experiment (until removal of the mussels) had a significant effect on the DNA copy number in each mesocosm experiment when all density treatments were analysed together (GLM, Z = −7.224, P < 0.0001, AIC = 912.95). There is still some overdispersion with this model, however the overdispersion parameter indicates this is not substantial (overdispersion statistic = 1.3, Residual Deviance 91.142 on 70 df, *χ^2^* P = 0.0457). There is substantial overlap between the DNA copy number estimates for the three density treatments (Fig. 2) and neither density nor total biomass significantly explained the number of DNA copies amplified by qPCR (Density Z = −1.019, P = 0.308; Total biomass Z = −0.776, P = 0.438). DNA was still detectable 24 hours after removal of the mussels, but not at 7 days after removal (Day 28) (Fig. 2). All blanks and mesocosm control tank samples were negative of target DNA throughout the experiment.

**Figure 2:**
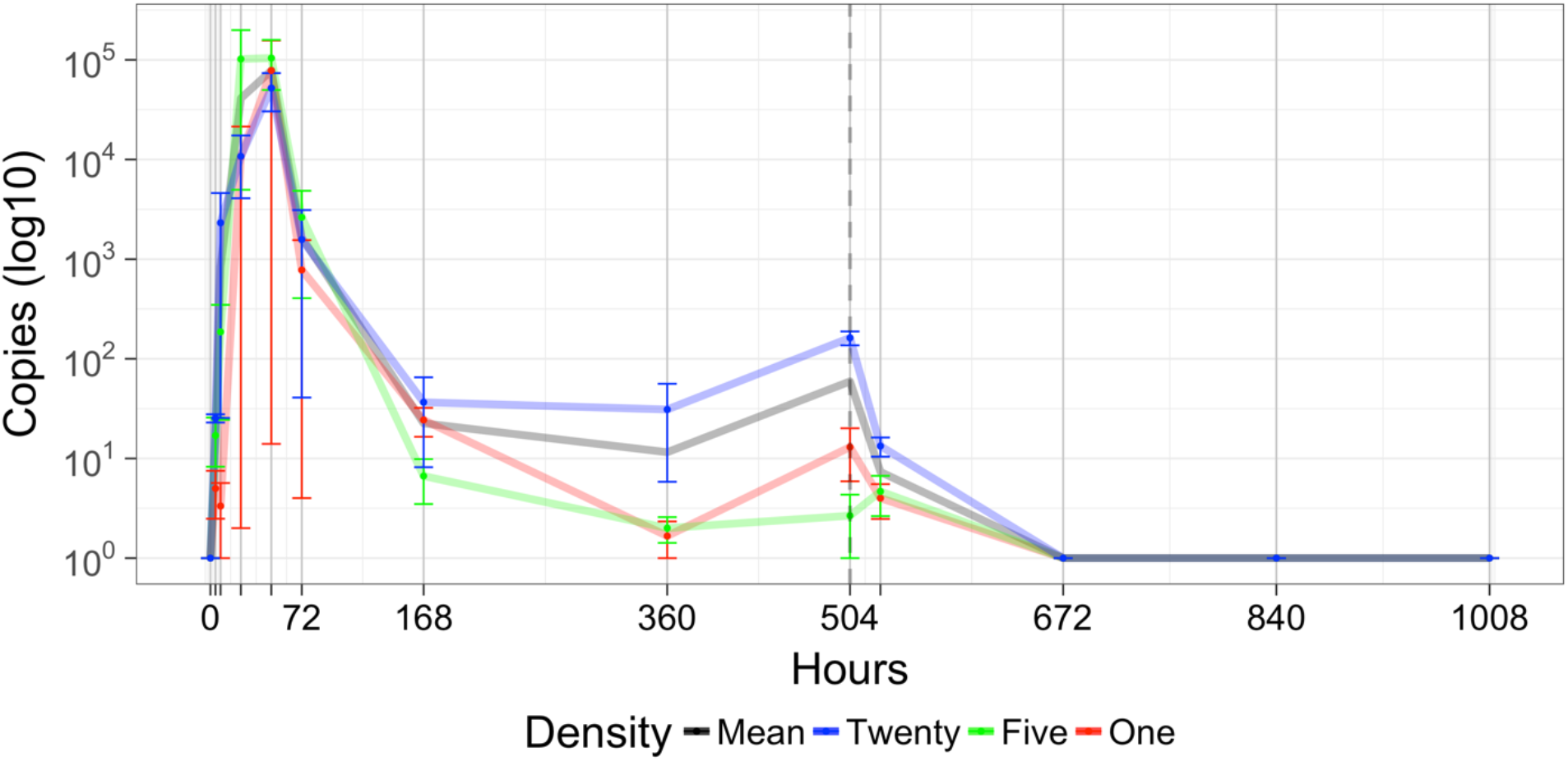
*Dreissena rostriformis bugensis* qPCR mesocosm assay validation. The graph shows the mean and standard deviation of DNA copy number recorded for each density treatment during the mesocosm experiment: red line – 1 specimen, green line – 5 specimens, blue line – 20 specimens and the black line is the mean copy number for all three densities. The vertical dashed line indicates when the mussels were removed from the mesocosm experiment.

### Detection of D. r. bugensis by traditional and molecular approaches in the field

#### Kick-net sampling

The number of *D. r. bugensis* specimens varied from 14 at Wraysbury Weir, the location closest to the reservoir outfall (source population), to a single specimen at Wraysbury Gardens, the location furthest from the outfall (Table 1). Detection rate was also highest at Wraysbury Weir (mussels detected in 4/6 sampling events). Mussels were detected in 1/6 samples at each of the other two sampling locations (Table 1).

**Table 1 A - C:**
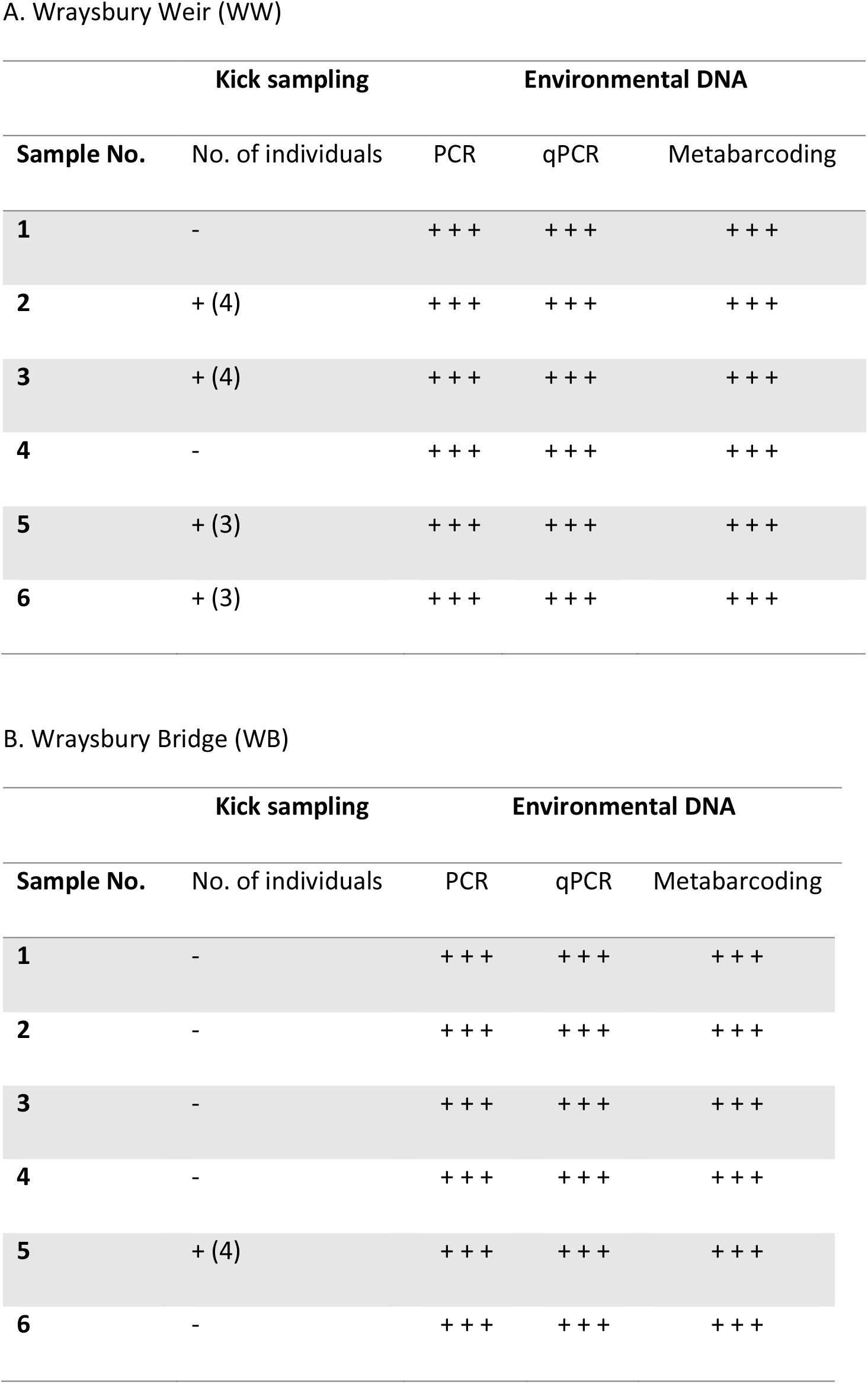
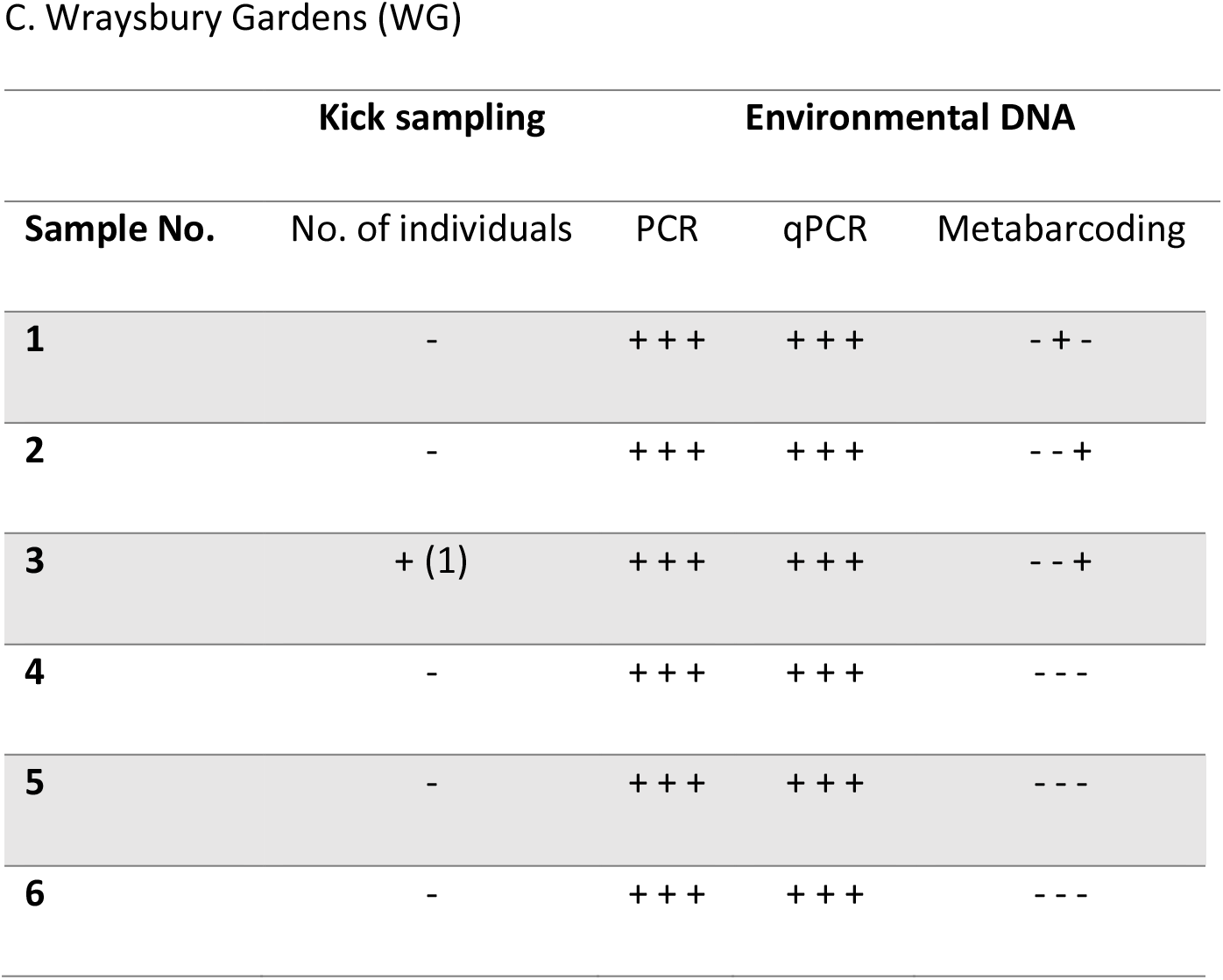
Comparison of methods for the detection of *D. r. bugensis* at three locations on the Wraysbury River. “+” indicates a positive detection and “−” indicates no detection for each technical replicate (i.e. three per sample for the eDNA methods and one for the kick sample). The numbers in parentheses under Kick Sampling correspond to the number of individuals found in each sample.

#### Targeted approaches

Detection rate was 100% in all three field locations and all replicates tested for both targeted approaches (cPCR and qPCR) (Table 1 and Fig. S3). Fig. 3A shows the DNA copy number (log_10_) of the qPCR assay for *D. r. bugensis* found at each of the sampling locations. Both distance from source and number of mussels found at the three locations, significantly influenced the DNA copy number, but distance was the strongest predictor (Density Z = −3.367, p = 0.000761; Distance Z = −17.223, p < 0.0001; AIC: 1392.1, Residual Deviance 57.621 on 51 df, χ^2^ P = 0.244). A Spearman’s Rank correlation between the DNA copy number and distance from the source population at each of the seven locations showed a weak correlation (ρ = −0.25, p = 0.5948). However, the low copy numbers found immediately downstream of the reservoir outfall (Fig. 3A) affected this. If this site is treated as an outlier and removed from the analysis, the correlation is significant (ρ = −1, p = 0.002778).

**Figure 3:**
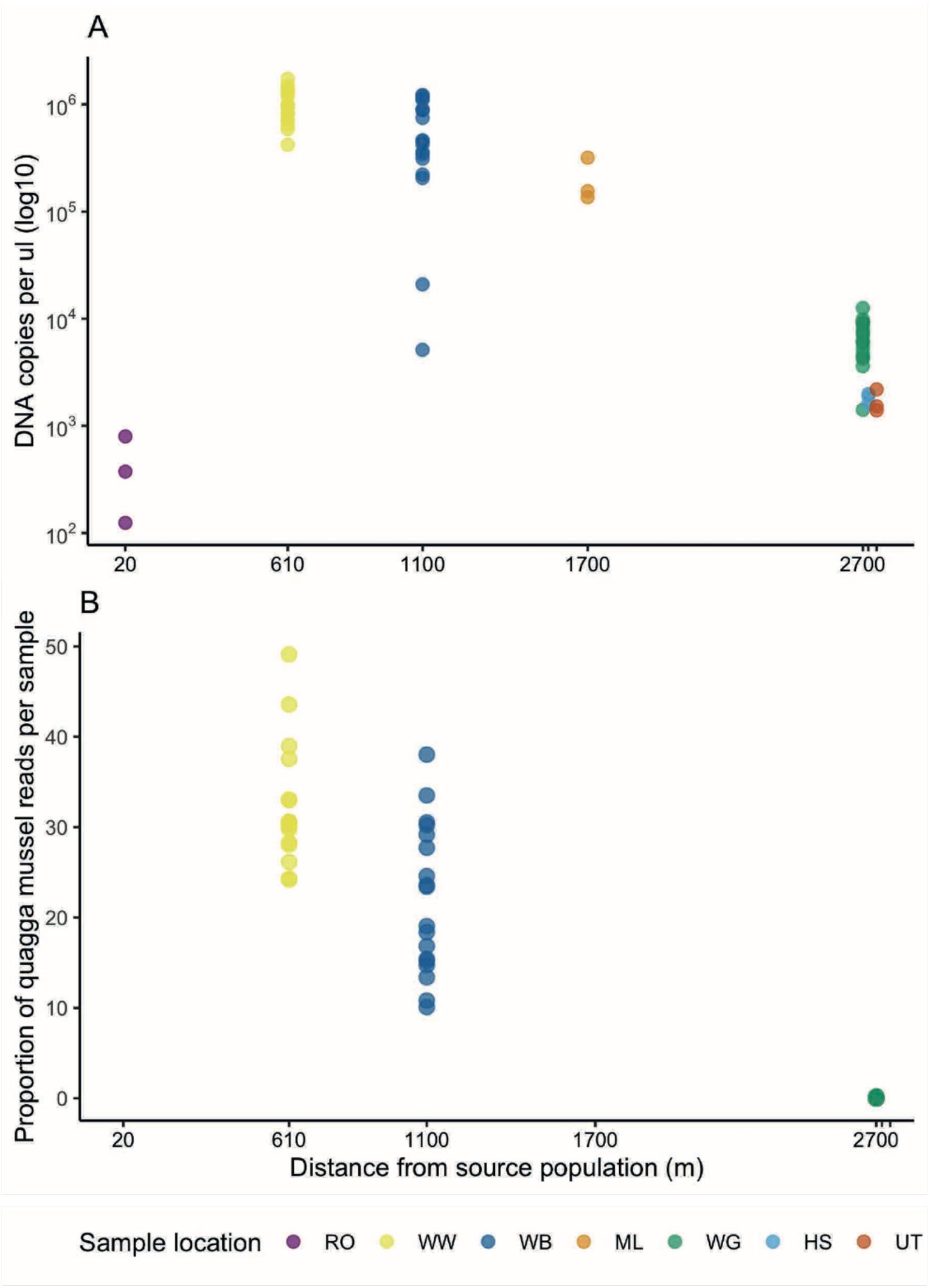
*Dreissena rostriformis bugensis* DNA signal recorded at field sites. A - qPCR DNA copy number per μl (log_10_) at all 7 sample sites. B – Normalised relative read count of taxa assigned to quagga mussel at the three locations sequenced.

#### Passive approach

The total number of reads produced following quality control and chimera removal was 4,928,265, of which 576,365 reads were assigned to the samples in this study. A total of 5.19% reads (29,936) were assigned to *D. r. bugensis*. After applying a 0.2% threshold to remove minor contamination the average read depth per sample was 2921 over all taxa, and 548 for *D. r. bugensis* (excluding controls). The clustering procedure identified 81 OTUs revealing a range of macroinvertebrate and algae/diatom species (Supplementary information III, Fig. S4). The proportion of reads assigned to *D. r. bugensis* from each sample within each location varied greatly as seen in Fig. 3B (n = 54). The target species was detected in all samples from both Wraysbury Weir and Wraysbury Bridge locations, while detection at Wraysbury Gardens was much lower with only 3 of the 18 replicates present for *D. r. bugensis* (Table 1). Detection rate was therefore 86%, and is lower than the targeted methods. The average raw read count of *D. r. bugensis* was 843 and 801 at Wraysbury Bridge and Wraysbury Weir respectively, and only 1 read on average at Wraysbury Gardens. Distance was the strongest predictor of the relative read count (Z = 13.47, p < 0.0001; AIC = 301.86, Residual Deviance 54.989 on 52 df, χ^2^ P = 0.36) in agreement with the qPCR results and when compared to the distance and density model (Distance-Z = −4.707, p < 0.0001; Density Z = −3.773, p = 0.0001; AIC = 258.64, Residual Deviance 49.696 on 51 df, χ^2^ P = 0.820), and density on its own (which was overdispersed).

## Discussion

This study investigates the detection, persistence and quantification of the invasive non-native quagga mussel (*D. r. bugensis*) using kick-net sampling and environmental DNA (eDNA). Through experimental mesocosms, we validated a species-specific DNA assay (cPCR) to detect quagga mussel and further developed a dye-based quantitative PCR (qPCR) assay to evaluate the performance of both targeted approaches. We carried out an eDNA survey of water samples and compared targeted (cPCR and qPCR) and passive (DNA metabarcoding) approaches to traditional kick-net sampling for qualitative and semi-quantitative detection in the River Wraysbury, UK. In line with our first hypothesis, detection rate was higher for all three eDNA methods than with the traditional method. Secondly, targeted approaches were more sensitive than the passive approach (100% detection rates versus 86%). Contrary to our third hypothesis, we observed no increased sensitivity using a dye-based qPCR over cPCR. Lastly, both density and distance from the source population were significant predictors of relative read count (metabarcoding) and copy number (qPCR) indicating that both approaches could provide estimates of quagga mussel abundance in the field.

### qPCR validation

Mesocosm validation provides an opportunity to test the LoD of eDNA assays and to investigate the relationship between density, time and DNA copy number (Dejean et al., 2011; Thomsen et al., 2012; Blackman et al., 2020). DNA was detected with qPCR at all three densities (1, 5 and 20 individuals) within 4 hours of being present, in agreement with previous cPCR results (Blackman et al., 2020). The LoD from serial dilutions of tissue DNA were also identical for cPCR and qPCR (1 x 10^−4^ ng/μl). Surprisingly, the mesocosm experiments highlighted unexpected dynamics of DNA in closed systems. Firstly, time since the start of the experiment was a significant predictor of DNA copy number, but rather than increasing over time or reaching a plateau, DNA accumulated quickly in the first 48 hours then rapidly depleted, and this trend was seen in all three density treatments. A possible explanation for this is stress induced on the mussel caused by a change in environment, as reported in other eDNA studies (Li et al., 2019). Dreissenid mussels, however, are known to “shut-down” in times of stress, such as chemical dosing, and can remain with their shells closed for up for three weeks (Aldridge et al., 2006). A further explanation for the rapid accumulation in DNA may be due to the production of protein based byssal threads which allow the mussels to attach to the substrate (Ricciardi et al., 1998; Aldridge et al., 2004; Timar and Phaneuf, 2009; Peyer et al., 2009). During the first 48 hours of the experiment, the mussels are likely to secure themselves to the new substrate. All mussels were found to have done this when they were removed at the end of the experiment. The mussels are also filter-feeders and were actively feeding (on microorganisms found in the river water) throughout the experiment producing sloughed cells, faeces and pseudofaeces (Sansom and Sassoubre, 2017). This feeding behaviour is likely to be a major contributor to successful detection in the field, and in a closed system, effect the DNA copy number detected with the qPCR assay.

The second surprising result from the mesocosm experiments was that we found no effect of density or biomass on DNA copy number. This contradicts several previous studies that have demonstrated a relationship between biomass or density and DNA copy number (Thomsen et al., 2012; Takahara., et al 2012; Doi et al., 2015). This could in part be due to the physiology and feeding behaviour of mussels, since eDNA is being consumed as well as produced (Friebertshauser et al., 2019). The complex pattern of eDNA accumulation and depletion together with the lack of discrimination between different density treatments presents challenges for estimation of quagga mussel abundance. However, eDNA is unlikely to accumulate in the same way in natural (particularly lotic) environments, and this case highlights that mesocosm experiments may poorly reflect the true relationship between DNA copy number and mussel density in the field. It would be worthwhile to examine whether this pattern is also observed with other filter feeding species. In keeping with previous studies (e.g. Thomsen et al., 2012), rapid degradation of eDNA was observed in mesocosm experiments, with a complete loss of signal by one week after mussel removal. Unfortunately, our experiments did not allow us to fully investigate the degradation rate of eDNA in mesocosms since our sampling was too limited. We recommend future studies carry out more intensive sampling once specimens have been removed.

### Comparison of eDNA and kick-net sampling for quagga mussel detection

The lower sensitivity of kick-net sampling is highlighted by the failure to collect a *D. r. bugensis* specimen in any of the first kick-net sampling attempts at each site. As demonstrated in a previous study, species in low abundance are likely to be missed, with only 62% of macroinvertebrate families detected in a three-minute kick-net sample (Furse et al., 1981). This highlights the need for multiple replicates for detection of low abundance species if using kick-net sampling. It should be noted that surveillance methods specifically for *D. r. bugensis* would be better focussed on the preferred habitat of mussels i.e. hard substrates such as man-made structures (Aldridge et al., 2014), which is more likely to detect quagga mussels than the randomised method used here. However, randomised kick-net sampling along a range of substrate types is the most widely used sampling method for freshwater macroinvertebrate monitoring, and therefore our experiment reflects the most likely methods to detect new INNS through routine monitoring schemes.

Of the three molecular approaches compared in this study, we show that both targeted approaches were more sensitive for detecting *D. r. bugensis* than metabarcoding, but that metabarcoding still performed encouragingly well, given that the assay targeted metazoan in general rather than the focal species. Few studies have compared targeted approaches with metabarcoding; however, our findings are similar to those of Simmons et al., (2015) and Harper et al., (2018), who showed that metabarcoding using conserved primers was less sensitive than targeted approaches. In this study, there was no increase in the detection of *D. r. bugensis* when the 0.2% contamination threshold was removed.

However, it would be pertinent to examine the raw output of metabarcoding runs before thresholds are applied in order to identify any potential INNS with low read counts. As this form of INNS monitoring is still in its infancy, quality control measures to ensure these detections are correct are yet to be put in place (Darling et al., 2020). We would therefore recommend any new INNS detections with metabarcoding are followed up with species-specific primers or collection of specimens prior to management actions to verify the detection.

A significant advantage of metabarcoding with conserved primers is the added information from non-target species detection: in our study a further 81 OTUs were identified, and although no additional INNS were detected, previous studies have demonstrated the power of this approach for detecting new and unexpected INNS (Blackman et al., 2017). Metabarcoding also provides the opportunity to monitor changes in community composition as the result of INNS or other ecological stressors (Simmons et al., 2015; Hänfling et al., 2016), which is extremely promising for better understanding INNS impacts for more effective prioritisation and management. This information on the wider community is a justification for the use of universal primers rather than more taxon-specific metabarcoding assays, but if more targeted information is needed, more specific metabarcoding primers can be developed. Indeed, specific mollusc metabarcoding primers have recently been developed, and are very promising for more targeted investigation of mollusc communities (Klymus et al., 2017). We hypothesize that more focussed metabarcoding assays will yield even higher detection rates than the one reported here, reducing the difference in sensitivity between targeted and passive approaches. It could also be possible to improve the power of metabarcoding by increasing spatial sampling. Future studies should investigate how many samples are needed in order to maximise the probability of detecting rare species with metabarcoding and also the required read depth needed per sample in order to maximise species detection at a given sampling site.

Conventional PCR and dye-based qPCR had identical detection rates for *D. r. bugensis* in our study. This is somewhat surprising as qPCR is generally considered to be more sensitive than cPCR (e.g. Thomsen et al., 2012, Nathan et al., 2014). Conventional PCR has the advantage of being quick and cost-effective, requiring limited lab equipment and up to half the time for sample processing than qPCR (Davison et al., 2016). Drawbacks to cPCR are typically lower sensitivity, and gel-quality issues that make interpretation of results more subjective than with qPCR. This is dependent on the primer specificity and optimisation of the PCR. Our conventional *D. r. bugensis* assay is highly specific and fully optimised, yielding clear bands on agarose gels even in samples with low concentration of target species DNA (as inferred from qPCR). However, we acknowledge that our experience with this assay may be unusual. An obvious advantage to qPCR is the quantitative information it provides, in addition to increased sensitivity. The added information on DNA concentration from qPCR in this study yielded important insights into the mussel density and distance from the source population (see below), which was not possible with cPCR. An important consideration is whether the assay could be improved by using probe-based rather than dye-based qPCR. Addition of a probe is generally considered beneficial to prevent the possibility of non-target amplification. Given the high specificity of the DRB1 primer pair described here, we believe the addition of a probe is not necessary in this case, but we stress that adding a probe should be considered if an extra layer of specificity is needed and/or if non-target amplification is possible.

### Quagga mussel eDNA declines with distance from the source and decreasing mussel density

Results from field trials for both dye-based qPCR and metabarcoding demonstrate a significant decrease in DNA copy number and relative read count of quagga mussel, respectively, with increasing distance from the main source population (Wraysbury reservoir) and with decreasing mussel density. Kick-net sample data also indicates that mussel density decreases along the course of the river, with only one mussel found at Wraysbury Gardens, 2.7 km downstream of the reservoir. The only site that did not fit this trend was the first site, 20 m downstream of the outfall from Wraysbury reservoir, which had a very low DNA copy number. The decrease in concentration of quagga mussel eDNA detected with distance from Wraysbury Reservoir, confirmed with both qPCR and metabarcoding, reflects both the decreasing density of mussels and a gradient in the transport of DNA downstream. The fact that eDNA concentration is not homogeneously distributed in the river suggests that it may be possible, at least in some cases, to infer the source population in cases where it is unknown. This is somewhat surprising given that our sampling took place after a period of heavy rain and eDNA distribution in rivers is often considered to be uniform and heavily confounded by flow rate and mixing (e.g Jane et al., 2015). Characteristics including discharge (Jane et al. 2015), physical structure and substrate type (Wilcox et al., 2016; Shogren et al., 2017) and water chemistry (Seymour et al., 2018) all influence the transport and retention of eDNA in lotic systems. Further empirical studies in rivers of all sizes and characteristics, ideally in combination with hydrological modelling (Carraro et al., 2018), are needed to fully understand the influence of flow dynamics and substrate type on retention and transport of eDNA in rivers.

Estimating abundance with eDNA is challenging because of the combined physical, chemical and biological factors that influence production, transport and retention of DNA in a water body (Barnes et al., 2014, Barnes and Turner 2015). Abundance estimation is expected to be particularly challenging in rivers due to their complex hydrodynamics, and the transport of eDNA from upstream (e.g. Jerde et al., 2016). Abundance estimation with metabarcoding is also controversial as read counts are influenced by bias during sampling, laboratory and bioinformatics steps (Valentini et al., 2016). Despite these challenges, our data add to the growing body of evidence that both qPCR and metabarcoding can yield insight into relative biomass, which has rarely been demonstrated in lotic environments (but see e.g. Doi et al., 2016, Akamatsu et al., 2020). Temporal sampling and controlling for environmental variables, combined with improved statistical approaches such as site occupancy modelling, which is becoming increasingly popular in eDNA studies (e.g. Schmidt et al., 2013, Pilliod et al., 2013, Chambert et al., 2018, Sales et al., 2020) or spatially explicit approaches that take hydrological conditions into account (e.g. Shogren et al., 2017, Carraro et al., 2018) will provide better quantitative information, and allow changes in biomass to be monitored.

### Conclusions and practical considerations for future INNS biomonitoring with eDNA

We have demonstrated that eDNA from quagga mussels can be successfully detected using cPCR, qPCR and metabarcoding in a UK river. Both qPCR and metabarcoding provide quantitative information, with DNA copy number and relative read count significantly decreasing with a combination of distance from the source population and mussel density. In our previous work we demonstrated that metabarcoding is an effective tool for passive detection of a previously unrecorded cryptic non-native species, *Gammarus fossarum* (Blackman et al. 2017). The present study provides further demonstration that metabarcoding with conserved metazoan primers is suitable as an early warning tool for non-targeted monitoring or routine surveillance even at a very low population density. The optimal approach for detection of INNS using eDNA will depend on the objectives of the research or monitoring programme. For early detection of previously unrecorded INNS, we recommend a comprehensive, randomised sampling design, combined with metabarcoding using universal primers, which can be supplemented with targeted approaches if additional verification is needed. Targeted qPCR assays provide optimal sensitivity and specificity and are particularly suited to surveys of INNS that have been previously recorded. In terms of future monitoring of quagga mussels, we believe that the sampling and laboratory methods adopted in the present study are widely applicable, but a temporal study would be beneficial to determine the optimum timing for sampling, and greater spatial coverage is needed to investigate the influence of environmental variables on detection rates.

## Acknowledgments

This project was funded by the Freshwater Biological Association Gilson Le Cren Award 2017 and the UK Environment Agency. We would like to thank staff from the Environment Agency - Drs Alice Hiley and Kerry Walsh for their advice and support and Dr Patrycja Meadows and Colette Sales for their help to identify field sites, arrange access and collect specimens. We also thank two anonymous reviewers for helpful and constructive suggestions on a previous version of our manuscript.

## Author contributions

The design and concept of this study was developed by RCB and LLH. RCB and LH carried out the field work. RCB, KKSL and PS performed the lab work. RCB analysed the data. RCB and LLH wrote the paper. All authors have approved the final manuscript.

## Data availability statement

Data generated in this study is available from the GitHub repository: https://github.com/RosettaBlackman/Blackman_et_al_quagga

