## Supplementary Information for "Targeted and passive environmental DNA approaches outperform established methods for detection of quagga mussels, *Dreissena rostriformis bugensis* in flowing water"

### Supplementary Material I – Methods

**Table S1: Sample sites used in this study**: Site information for the points where kick-net and eDNA samples were collected for each experiment on the River Wraysbury. Note: those eDNA samples analysed with both methods (i.e. qPCR and metbarcoding) are not collected in duplicate but were just analysed with the two methods.

| Sample type | Site Name | Grid ref | Kick-net samples | eDNA samples – qPCR analysis | eDNA samples – metabarcoding analysis |
|---|---|---|---|---|---|
| **Field samples** | Wraysbury reservoir outfall | TQ0304374214 | N/A | 1* 3 | N/A |
| | Wraysbury Weir | TQ0289873636 | 6 | 6* 3 | 6* 3 |
| | Wraysbury Bridge | TQ0267973201 | 6 | 6* 3 | 6* 3 |
| | Moor Lane | TQ0280272634 | N/A | 1* 3 | N/A |
| | Wraysbury Gardens | TQ0334071906 | 6 | 6* 3 | 6* 3 |
| | Hale Street | TQ033371745 | N/A | 1* 3 | N/A |
| | Upstream Thames confluence | TQ0334671701 | N/A | 1* 3 | N/A |
| | | **Subtotal** | **18** | **66** | **54** |
| **Blanks controls** | Field Blank (Waterside Car Park) | TQ0487177572 | N/A | 1* 3 | 1*3 |
| | Filter Blank | N/A | | 1* 3 | 1* 3 |
| | | **Total** | **18** | **72** | **60** |

*Water sampling and DNA extraction*

Water filtration was carried out in the dedicated eDNA facility at the University of Hull, UK. Five-hundred millilitres of each sample was filtered and extracted separately to allow replication within each eDNA sampling event. Samples were vacuum filtered through sterile 47 mm diameter 0.45 µm cellulose nitrate membrane filters with pads (Whatman, GE Healthcare, UK) within 24 hours of collection, using Nalgene filtration units (Thermo Fisher Scientific) in combination with a vacuum pump (15~20 in. Hg, Pall Corporation). All sampling and filtration equipment was sterilized in 10% commercial bleach solution for 10 minutes then rinsed with 10% MicroSol detergent (Anachem, UK) and purified water between each sample. All DNA extractions in this study were carried out using the method by Brolaski et al., (2008) with minor modifications (See Supplementary Information III).

*Conventional PCR protocol*

Conventional PCRs were carried out in triplicate in 25 µl volumes with 2x MyTaq Red Mix Taq (Bioline, UK) containing: 0.4 µM of each primer, and 2 µl of undiluted DNA template. PCRs were performed on an Applied Biosystems Veriti Thermal Cycler with the following profile: initial denaturation at 94°C for 3 min, followed by 37 cycles of denaturation at 94°C for 30s, annealing at 65°C for 1 min and extension at 72°C for 1 min 30s, with a final extension time of 10 min at 72°C. Detection was then confirmed by running PCR products on a 1.5% TBE agarose gel stained with GelRed (Cambridge Biosciences, UK). Our detection criteria required amplification of at least 2 out of 3 replicates.

*qPCR protocol*

Quantitative PCR reactions were carried out in triplicate on a StepOne-Plus™ Real-Time PCR machine (Applied Biosystems) in 25 µl reaction volumes. Primers (DRB1) and reagent concentration were the same for qPCR as for the conventional PCR reaction; with the replacement of MyTaq for Power Up SYBR Master Mix (Fisher Scientific, UK) in the qPCR reaction. qPCRs were performed with the following profile: 2 minutes at 50°C, 10 minutes at 95°C, and 50 cycles of 15 s at 95°C and 1 minute at 60°C followed by melt curve analysis. A 300 bp laboratory synthesized gBlocks® fragment (Integrated DNA Technologies Inc.) of the *D. r. bugensis* COI gene was used to construct a calibrated standard curve. DNA copy number for the gBlocks® fragment was estimated using Avogadro's number (number of copies = (500 ng of DNA × 6.022 × 1023) / (300 bp fragment × 1 × 109 × 650). This formula assumes that the average weight of a base pair is 650 Da. The lypholised gBlock was re-suspended in 50 µl of sterile 1x TE buffer, which made a stock concentration of 5.0 x $10^{10}$ copies. From this stock solution, one five-fold dilution followed by eight 10-fold dilutions were made to produce the standard curve (10 to $10^8$ copies per uL) which was used to calculate copy number in the samples.

*Metabarcoding workflow*

First step PCRs were performed with the primers jgHCO2198: 5'-TAIACYTCIGGRTGICC RAARAAYCA-3' and mICOIintF: 5'-GGWACWGGWTGAACWGTWTAYCCYCC-3' (Geller et al., 2013; Leray et al., 2013). PCRs were carried out in 50µl reaction volumes with Sigma Taq (Sigma Aldrich, UK) containing: 0.4 µM of each primer, 1.5 mM $MgCl_2$, 2 µM dNTPs, and 2 µl of undiluted DNA template. PCRs were performed on an Applied Biosystems Veriti Thermal Cycler machine with the following profile: initial denaturation 95°C for 2 mins, followed by 35 cycles of denaturation 95°C for 15s, annealing at 50°C for 30s and extension at 72°C for 30s, with a final extension time of 10 min at 72°C. All PCRs included PCR blanks and positive controls of DNA from species that are terrestrial (*Osmia bicornis*, n=1) or not found in the sampled water bodies (*Triops cancriformis*, n=2) were included on all PCR plates. PCR products were confirmed by gel electrophoresis on a 1.5% TBE agarose gel stained with GelRed (Cambridge Bioscience, UK). PCR clean-up was carried out using ZR-96 DNA Clean-up Kit™ (Zymo Research, USA) following the manufacturer's protocol.

Individual Nextera tags (Illumina, UK) were then added in a second PCR with 10 µM of each tagging primer and 1 µl of purified PCR product. PCR settings were: initial denaturation 95°C for 2 min, followed by 8 cycles of denaturation 95°C for 15s, annealing at 55°C for 30 secs and extension at 72°C for 30 secs, with a final extension time of 10 mins at 72°C. Samples were then normalised using SequalPrep Normalization plates (Invitrogen, UK). Each plate of samples was then pooled and the library made with equimolar concentrations of each plate. The final pooled library was concentrated by SpeedVac vacuum and then cleaned using the QIAquick Gel Extraction Kit (Qiagen, UK) following the manufacturer's protocol. The library was sequences at 13 pM concentration with 20% PhiX, on an Illumina MiSeq platform using 2 x 250bp v2 chemistry at the University of Hull.

Processing of Illumina read data and taxonomic assignment were performed using a custom bioinformatics pipeline MetaBEAT (metaBarcoding and eDNA Analysis Tool) v 0.8 (https://github.com/HullUnibioinformatics/metaBEAT). The program Trimmomatic 0.32 (Bolger et al., 2014) was used for quality trimming and removal of adapter sequences from the raw Illumina reads. Average read quality was assessed in 5 bp sliding windows starting from the 3'-end of the read and reads were clipped until the average quality per window was above phred 30. Sequence pairs were subsequently merged into single high-quality reads using the program FLASH 1.2.11 (Magoč & Salzberg., 2011). All paired reads shorter than a defined minimum read length (313bp) were discarded. The remaining reads were screened for chimeric sequences against a reference database using the 'uchime_ref' function implemented in vsearch 1.1 (Rognes et al. 2016). To remove redundancy, sequences were clustered at 100% identity using vsearch 1.1. Clusters represented by less than 5 sequences were considered sequencing error and were omitted from further analyses. Non-redundant sets of query sequences were then compared to full the GenBank database using BLAST (Zhang et al., 2000). BLAST output was interpreted using a custom python function, which implements a lowest common ancestor (LCA) approach for taxonomic assignment similar to the strategy used by MEGAN (Huson et al., 2007). In brief, after the BLAST search we recorded the most significant matches to the reference database (yielding the top 10% bit-scores) for each of the query sequences. If only a single taxon was present in the top 10%, the query was assigned directly to this taxon. If more than one reference taxon was present in the top 10%, the query was assigned to the lowest taxonomic level that was shared by all taxa in the list of most significant hits for this query. Sequences for which the best BLAST hit had less than 97% identity to any sequence on GenBank, were considered non-target sequences and discarded. Sequences were then filtered for the presence of *D. r. bugensis* using GenBank sequences: EF080862, JX945980, EF080861, U47651, AF495877, EU484436, KJ881409. We quantified the level of contamination by examining the species detected in the single species positive samples (*Osmia bicornis* and *Triops cancriformis*), 42 reads assigned to Ascomycota in Positive 10.  Low level contamination was found from human (5 and 14 reads) and positive tissue samples in the negative controls (*T. cancriformis* 10 reads). This was addressed by applying a 0.2% contamination threshold, i.e. removing sequences that had a frequency of up to 0.2% of the total reads in each sample.

### Supplementary Information III - DNA Extraction method

**Protocol for DNA extraction from filter papers modified from Brolaski et al., (2008)**

Lysis solution 1 - 0.12µM Guanidine thiocyanate and 0.181 µM Tridocium phosphate

Lysis solution 2 – 5 µM sodium chloride, 0.5 µM Tris base, 4% SDS

Precipitation solution – 5 µM ammonium acetate, 0.12 alluminium ammonium sulphate dodecahydrate

Binding solution – 5 µM Guanidine HCl, 0.03 µM Tris HCl, 9% Isoproponol

Wash solution – 0.01 µM Tris HCl, 0.5 µM Sodium Chloride, 75% Ethanol

Elution Buffer – TE buffer

1. 1g 30mesh garnet beads, 1g fine sand into 7ml tube
2. Add filter paper
3. 925 µl Lysis solution 1 and 75 µl Lysis solution 2
4. Qiagen Tissue lyser 5 minutes, 30 bps
5. Centrifuge 4000g, 1min
6. Pipette off supernatant into clean 2ml tube
7. Add 250 µl Precipitation solution, vortex
8. Chill on ice for 5 mins
9. Centrifuge 10000g, 1min
10. Pipette off supernatant into clean 2ml tube
11. Add x1.5 volume of Binding solution, vortex
12. Pipette 650 µl into spin column, centrifuge 10000g, 1 min, discard flow through
13. Repeat step 12 until all solution has gone through spin column

14. Add 500 µl l of Wash solution, centrifuge 10000g, 1 min, discard flow through
15. Centrifuge spin column 10000g, 2 min
16. Place spin column in fresh collection tube
17. Add 100 µl of Elution buffer (TE, dd$H_2O$) leave for 5 minutes
18. Centrifuge 10000g, 1 min.

### Supplementary Information III – Tables and Figures

Table S2: GLM analyses of DNA copies for quagga mussel produced by qPCR of field samples

Table S3: GLM analyses of DNA copies for quagga mussel produced by qPCR of mesocosm samples.

Table S4: GLM analyses of metabarcoding reads for quagga mussel in field samples.

Figure S1: qPCR product visualised on an agarose gel to show amplification.

Figure S2. Agarose gel images from *D. r. bugensis* field samples collected from River Wraysbury.

Figure S3: qPCR (SYBR green) amplification plot (a) and melt curve (b) for quagga mussel, *Dreissena rostriformis bugensis*.

Figure S4: Other taxa detected from the metabarcoding analysis.

**Table S2: GLM analyses of DNA copies for quagga mussel produced by qPCR of mesocosm samples.** This table gives the output from GLMs performed in R (R-Core-Team 2017)

| Model | AIC | Estimate Std. Error | z value | Pr(>\|z\|) (p-value) | Residual deviance | Residual deviance P-val (1-pchisq)* |
|---|---|---|---|---|---|---|
| Hours + TotalBio | 914.94 | 0.002236<br>0.044271 | -7.265<br>0.078 | 3.73e-13 ***<br>0.938 | 91.141 on 69 df | 0.03840271 |
| Hours + Density | 914.87 | 0.002233<br>0.042593 | -7.055<br>-0.354 | 1.72e-12 ***<br>0.723 | 91.138 on 69 df | 0.03842042 |
| TotalBio | 925.27 | 0.04715 | -0.776 | 0.438 | 92.236 on 70 df | 0.03873005 |
| Hours | 912.95 | 0.002235 | -7.224 | 5.07e-13 *** | 91.142 on 70 df | 0.04567788 |
| Density | 924.91 | 0.04530 | -1.019 | 0.308 | 92.205 on 70 df | 0.03891364 |

**Table S3: GLM analyses of DNA copies for quagga mussel produced by qPCR of field samples.** This table gives the output from GLMs performed in R (R-Core-Team 2017)

| Model | AIC | Estimate Std. Error | z value | Pr(>\|z\|) (p-value) | Residual deviance | Residual deviance P-val (1-pchisq)* |
|---|---|---|---|---|---|---|
| Distance + Density | 1392.1 | 0.000173<br>0.027777 | -17.223<br>-3.367 | < 2e-16 ***<br>0.000761 *** | 57.621 on 51 df | 0.2436637 |
| Distance | 1400.7 | 0.0001063 | -23.78 | <2e-16 *** | 58.276 on 52 df | 0.2555702 |
| Density | 1491.1 | 0.03279 | 5.102 | 3.37e-07 *** | 67.462 on 52 df | 0.07331911 |

**Table S4: GLM analyses of metabarcoding reads for quagga mussel in field samples.** This table gives the output from GLMs performed in R (R-Core-Team 2017).

| Model | AIC | Estimate Std. Error | z value | Pr(>\|z\|) (p-value) | Residual deviance | Residual deviance P-val (1-pchisq)* |
|---|---|---|---|---|---|---|
| Distance+<br>Density | 258.64 | 0.0009511<br>0.0480304 | -4.707<br>-3.773 | < 2.51e-06 ***<br>0.0001 *** | 41.696 on 51 df | 0.820419 |
| Distance | 301.86 | 0.0001565 | -13.47 | <2e-16 *** | 54.989 on 52 df | 0.3621245 |
| Density | 694.99 | 0.005852 | 18.99 | <2e-16 *** | 505.52 on 52 df | 0 - overdispersed |

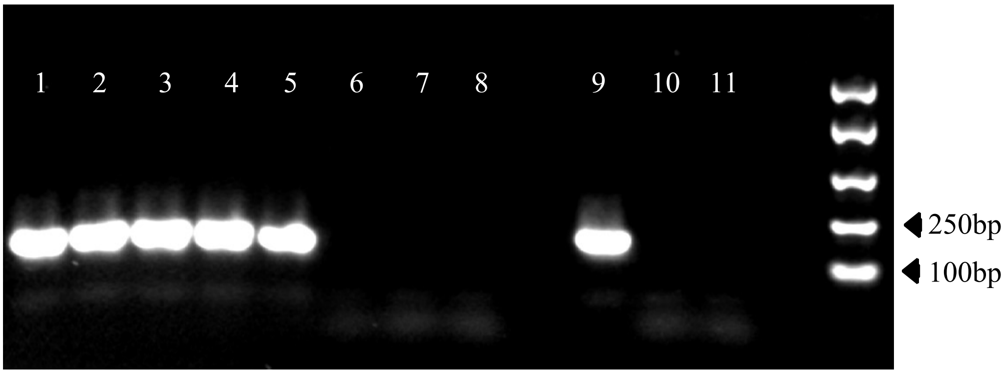


**Figure S1: qPCR product of serial quagga mussel DNA dilutions visualised on an 1.5% agarose gel to show amplification.** Lane 1: 1ng/µl, Lane 2: 1:10 dilution, 0.1 ng/µl per reaction, Lane 3: 1:100 dilution, 1 x $10^{-2}$ ng/µl per reaction, Lane 4: 1:1000, ~1 x $10^{-3}$ ng/µl per reaction, Lane 5: 1:10000, ~1 x $10^{-4}$ ng/µl per reaction, Lane 6: 1:100000, ~01 x $10^{-5}$ ng/µl per reaction, Lane 7: 1:1000000, ~1 x $10^{-6}$ ng/µl per reaction, Lane 8: 1:10000000, ~01 x $10^{-7}$ ng/µl per reaction, Lane 9 is a positive tissue sample and 10 and 11 are PCR negative ($ddH_2O$). Concentrations were measured on QuBit 2.0 prior to qPCR reaction and therefore Lanes 4-8 given are given as approximations only. The final lane is DNA EasyLadder I (Bioline, UK).

A. B.

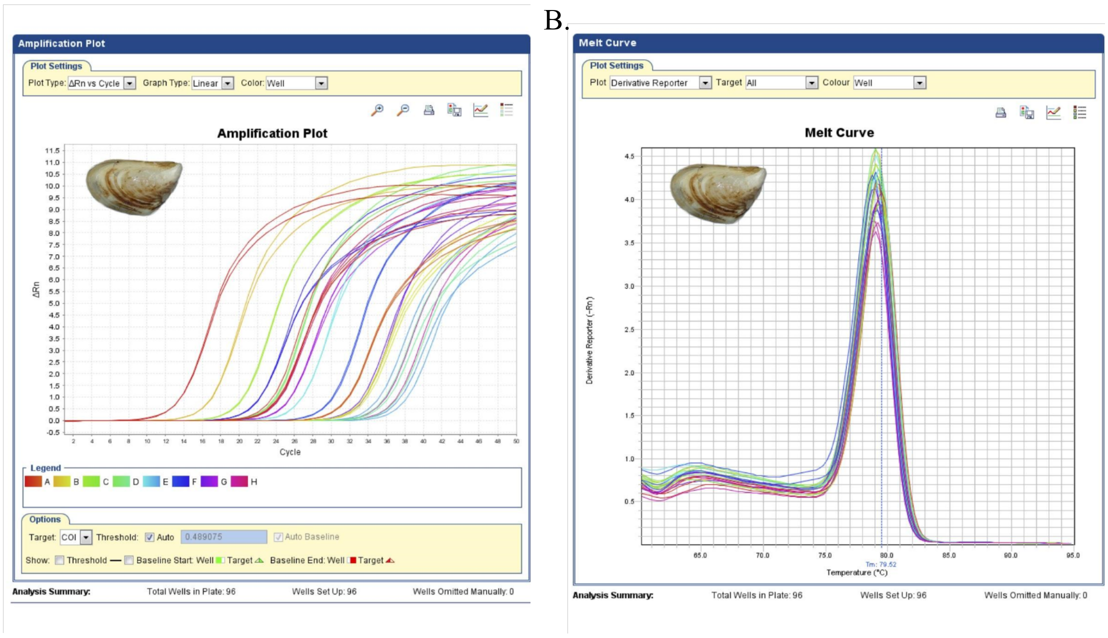


**Figure S2: qPCR (SYBR green) amplification plot (A) and melt curve (B) for quagga mussel, *Dreissena rostriformis bugensis*.** This example amplification plot represents the accumulation of PCR product over the duration of the qPCR experiment. A) The Y axis is the change in relative fluorescence and PCR cycle number on the X. Each coloured line on the plot represents a sample (in triplicate). The earlier in the qPCR experiment the curve increases from the baseline fluorescence (X axis), the greater the number of target DNA copies in the sample. B) The melt curve charts the change in fluorescence observed when double stranded DNA separates into single strands. Plotting the curve is a way of checking for reaction specificity. A single, distinct peak (as shown here) demonstrates a highly specific reaction.

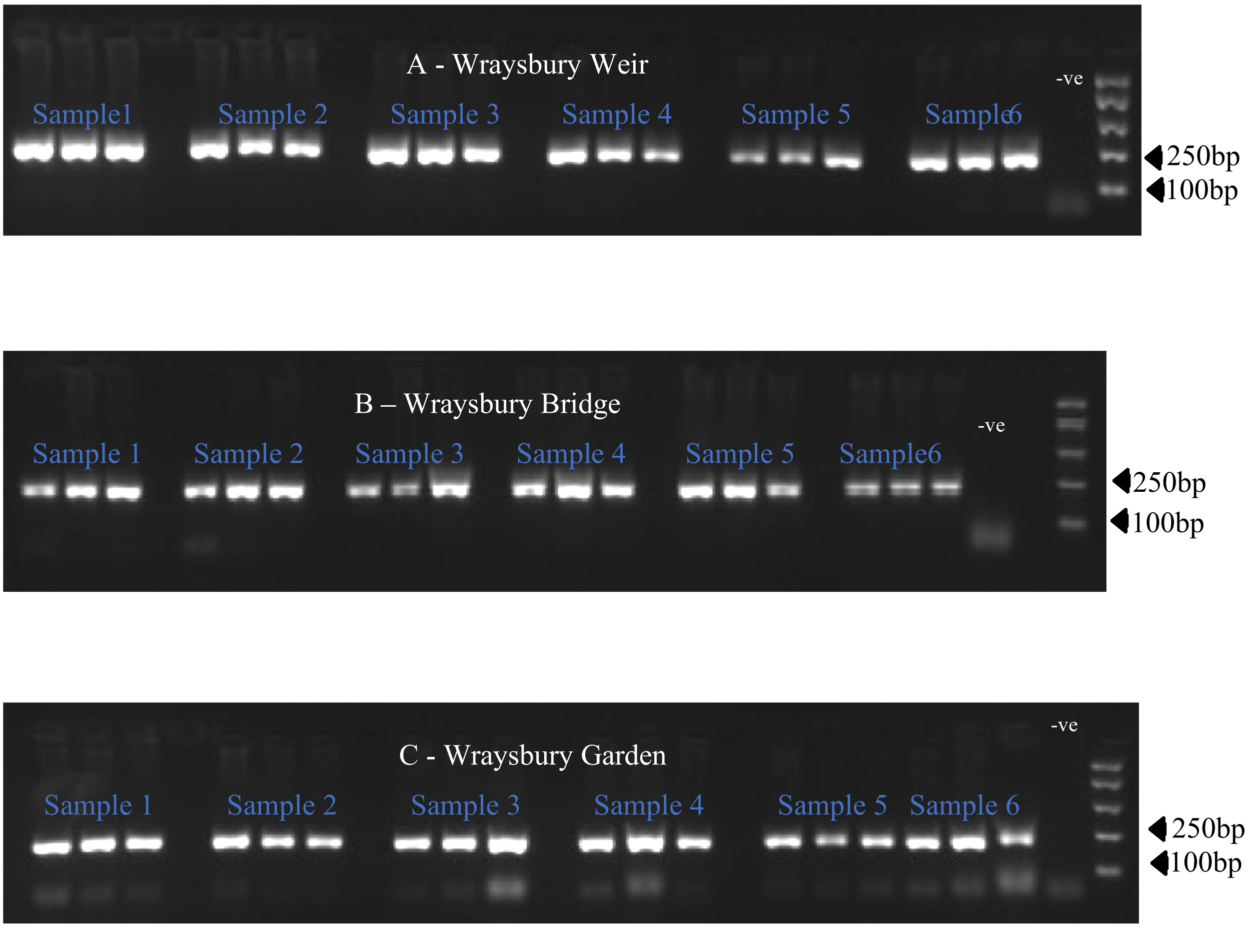


**Figure S3. 1.5% Agarose gel images from *D. r. bugensis* field samples collected from River Wraysbury.** A – Wraysbury Weir, B – Wraysbury Bridge and C – Wraysbury Gardens. For a detection to be deemed positive a band must be present at the correct size for a minimum of 2 out of the 3 replicates, as these gel images show all samples fulfilled these criteria. The final lane is DNA EasyLadder I (Bioline, UK) with corresponding fragment size.

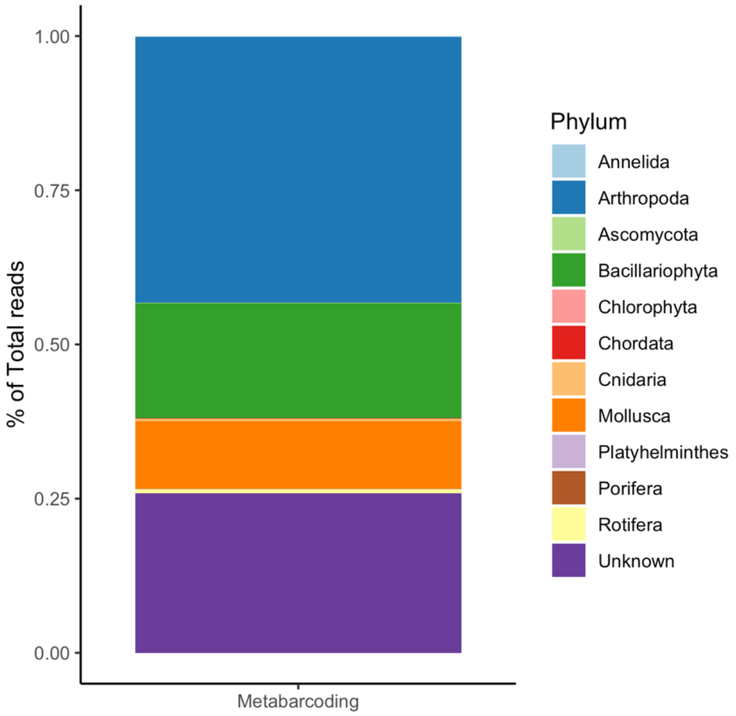


**Figure S4: Other taxa detected from the metabarcoding analysis.** A further 81 taxa were detected using the jgHCO2198 and mICOIintF primers set (Leray et al., (2013) and Geller et al., 2013), excluding the target species *Dreissena rostriformis bugensis*. This stacked bar chart shows the taxa to Phylum level to indicate the range of taxa detected using the metabarcoding approach (Unknown Phylum refers to taxa which are not classified at Phylum level from Classes Chrysophyceae, Dinophyceae, Florideophyceae, Oomycetes.)

#### Supplementary Information IV – MIQE Checklist

| *ITEM TO CHECK* | *IMPORTANCE* | *CHECKLIST* |
|---|---|---|
| EXPERIMENTAL DESIGN | | |
| Definition of experimental and control groups | E | Mesocosm experiment: 3 densities of *D. r. bugensis* (1, 5, and 20) with a tank with no specimens (control)<br>Field experiment: 7 sites sampled on the R. Wraysbury with a control sample upstream of *D. r. bugensis* spread |
| Number within each group | E | Mesocosm control: n=1, Experimental: n=9, Total: n = 10<br>Field experiment control: n = 3, Experimental: n = 66 |
| Assay carried out by core lab or investigator's lab? | D | Investigator's Lab |
| Acknowledgement of authors' contributions | D | Yes |
| SAMPLE | | |
| Description | E | eDNA water sample |
| Volume/mass of sample processed | D | 25ul |

| | | |
|---|---|---|
| Microdissection or macrodissection | E | N/A |

| | | |
|---|---|---|
| Processing procedure | E | Mesocosm: 200ml water samples were collected from each mesocosm at 4hrs, 8hrs, 24hrs, 7 days, 15 days, 21 days, 22 days, 28 days, and 42 days. Samples were vacuum filtered through sterile 47 mm diameter 0.45 µm cellulose nitrate membrane filters with pads (Whatman, GE Healthcare, UK) immediately after collection, using Nalgene filtration units (Thermo Fisher Scientific) in combination with a vacuum pump (15~20 in. Hg, Pall Corporation). Filtered samples were returned to the correct mesocosm_tank<br>Field: 3 x 500ml water samples were collected at each sampling point and processed within 24 hours in the same way as the mesocosm samples. All samples were stored in petri dishes at –20 °C until DNA extraction. DNA extractions were carried out using a protocol modified |

| | | |
|---|---|---|
| | | from Bolaski et al., (2008) (for the full extraction protocol, see Supplementary Information I). DNA was eluted with 100 µl of Buffer AE. The DNA solution was stored in a 0.8ml microtube at –20 °C until PCR analysis. |
| If frozen - how and how quickly? | E | N/A |

| | | |
|---|---|---|
| If fixed - with what, how quickly? | E | N/A |
| Sample storage conditions and duration (especially for FFPE samples) | E | All concentrated samples were stored in 0.8ml microtubes at –20°C. |
| NUCLEIC ACID EXTRACTION | | |
| Procedure and/or instrumentation | E | We used a modified Bolaski et al., (2008) protol as documented in Supplementary information I |
| Name of kit and details of any modifications | E | We used a modified Bolaski et al., (2008) protol as documented in Supplementary information I |
| Source of additional reagents used | D | University of Hull |
| Details of DNase or RNAse treatment | E | N/A |
| Contamination assessment (DNA or RNA) | E | N/A |
| Nucleic acid quantification | E | Quantification was performed using a Qubit 2.0 following the manufacturer's instructions. |
| Instrument and method | E | |
| Purity (A260/A280) | D | |
| Yield | D | |

| | | |
|---|---|---|
| RNA integrity method/instrument | E | N/A |
| RIN/RQI or Cq of 3' and 5' transcripts | E | N/A |
| Electrophoresis traces | D | N/A |
| Inhibition testing (Cq dilutions, spike or other) | E | N/A |
| REVERSE TRANSCRIPTION | | |
| Complete reaction conditions | E | N/A |
| Amount of RNA and reaction volume | E | N/A |
| Priming oligonucleotide (if using GSP) and concentration | E | N/A |
| Reverse transcriptase and concentration | E | N/A |
| Temperature and time | E | N/A |
| Manufacturer of reagents and catalogue numbers | D | N/A |
| Cqs with and without RT | D | N/A |

| | | |
|---|---|---|
| Storage conditions of cDNA | D | N/A |

| | | |
|---|---|---|
| qPCR TARGET INFORMATION | | |
| If multiplex, efficiency and LOD of each assay. | E | N/A |
| Sequence accession number | E | DQ840132.1 |
| Location of amplicon | D | amplicon location: 196 - 384 |
| Amplicon length | E | Including primers - 188 bp |
| In silico specificity screen (BLAST, etc) | E | The in-silico specificity screen was performed using Primer-BLAST. The *D. r. bugensis* primer pair, DRB1, amplified 29 published *D. rostriformis, D. bugensis* and *D. rostriformis bugensis* sequences in silico with no mismatches. |
| Pseudogenes, retropseudogenes or other homologs? | D | Not Found |
| Sequence alignment | D | N/A |
| Secondary structure analysis of amplicon | D | Not Checked |

| | | |
|---|---|---|
| Location of each primer by exon or intron (if applicable) | E | N/A |
| What splice variants are targeted? | E | N/A |
| qPCR OLIGONUCLEOTIDES | | |
| Primer sequences | E | DRB1_F(5'-GGAAACTGGTTGGTCCCGAT-3')<br>DRB1_R (5'-GGCCCTGAATGCCCCATAAT-3’) |
| RTPrimerDB Identification Number | D | Not Submitted |
| Probe sequences | D | N/A |
| Location and identity of any modifications | E | |
| Manufacturer of oligonucleotides | D | IDT Ltd |
| Purification method | D | HPLC |
| qPCR PROTOCOL | | |
| Complete reaction conditions | E | Reactions were set up manually in specialist eDNA laboratory. |

| | | |
|---|---|---|
| Reaction volume and amount of cDNA/DNA | E | Reaction volume is 25 µl, and amount of DNA is 2 µL |
| Primer, (probe), Mg++ and dNTP concentrations | E | N/A |
| Polymerase identity and concentration | E | We used Power Up SYBR Master Mix following manufacturer's instructions |
| Buffer/kit identity and manufacturer | E | We used Power Up SYBR Master Mix following manufacturer's instructions |
| Exact chemical constitution of the buffer | D | N/A |
| Additives (SYBR Green I, DMSO, etc.) | E | 12.5 ul |
| Manufacturer of plates/tubes and catalog number | D | Microamp, Optical 96 Well Reaction Plate (10411785) & Microamp Optical Adhesive Film (10299204)<br>(Applied Biosystems) |

| | | |
|---|---|---|
| Complete thermocycling parameters | E | 2 min at 50°C, 10 min at 95°C, and 55 cycles of 15 s at 95°C and 60 s at 60°C. |
| Reaction setup (manual/robotic) | D | We performed following the manufacturer's instructions. |
| Manufacturer of qPCR instrument | E | StepOne-Plus™ Real-Time PCR system (Applied Biosystems, Foster City, CA, USA) |
| qPCR VALIDATION | | |

| | | |
|---|---|---|
| Evidence of optimisation (from gradients) | D | Not Performed |
| Specificity (gel, sequence, melt, or digest) | E | The specificity of the primer was tested by sequencing PCR product by Macrogen |
| For SYBR Green I, Cq of the NTC | E | None |
| Standard curves with slope and y-intercept | E | Slope: Range -2.97 – -3.69, y-intercept: Range - 32.85–42.96 |
| PCR efficiency calculated from slope | E | 86.6–116% |
| Confidence interval for PCR efficiency or standard error | D | Standard error for PCR effciency = 1.68 |

| | | |
|---|---|---|
| r2 of standard curve | E | 0.9906–0.99969 |
| Linear dynamic range | E | Linear dynamic range from 124 to 1733209 DNA copies per reaction |
| Cq variation at lower limit | E | The positive signals were detected from the two of three wells for five copy template. |
| Confidence intervals throughout range | D | Not Checked |
| Evidence for limit of detection | E | Because five copies of target DNA was detected in at least two wells in each set of triplicates, we defined the limit of detection as 5 copies. |
| If multiplex, efficiency and LOD of each assay. | E | N/A |
| DATA ANALYSIS | | |
| qPCR analysis program (source, version) | E | StepOne Software ver 2.0 |
| Cq method determination | E | We performed according to default setting of Software above. |
| Outlier identification and disposition | E | |
| Results of NTCs | E | Three wells of no-template negative control were included in all qPCR plates and showed no amplification. |

| | | |
|---|---|---|
| Justification of number and choice of reference genes | E | N/A |
| Description of normalisation method | E | We used standard curve methods. |
| Number and concordance of biological replicates | D | N/A |
| Number and stage (RT or qPCR) of technical replicates | E | Triplicate for each qPCR. |
| Repeatability (intra-assay variation) | E | N/A |
| Reproducibility (inter-assay variation, %CV) | D | N/A |
| Power analysis | D | N/A |
| Statistical methods for result significance | E | We performed according to default setting of Software above. |
| Software (source, version) | E | StepOne Software ver 2.0 |
| Cq or raw data submission using RDML | D | Not Submitted |
